# Auditory and visual gratings elicit distinct gamma responses

**DOI:** 10.1101/2024.01.05.574448

**Authors:** Divya Gulati, Supratim Ray

**Affiliations:** Centre for Neuroscience, Indian Institute of Science, Bengaluru, India, 560012

## Abstract

Sensory stimulation is often accompanied by fluctuations at high frequencies (>30Hz) in brain signals. These could be “narrowband” oscillations in the gamma band (30-70 Hz) or non-oscillatory “broadband” high-gamma (70-150 Hz) activity. Narrowband gamma oscillations, which are induced by presenting some visual stimuli such as gratings and have been shown to weaken with healthy aging and the onset of Alzheimer’s Disease, hold promise as potential biomarkers. However, since delivering visual stimuli is cumbersome as it requires head stabilization for eye tracking, an equivalent auditory paradigm could be useful. Although simple auditory stimuli have been shown to produce high-gamma activity, whether specific auditory stimuli can also produce narrowband gamma oscillations is unknown. We tested whether auditory ripple stimuli, which are considered an analogue to visual gratings, could elicit narrowband oscillations in auditory areas. We recorded 64-channel EEG from male and female (18 each) subjects while they either passively fixated on the monitor while viewing static visual gratings, or listened to stationary and moving ripples, played using loudspeakers, with their eyes open or closed. We found that while visual gratings induced narrowband gamma oscillations with suppression in the alpha band (8-12Hz), auditory ripples did not produce narrowband gamma but instead elicited very strong broadband high-gamma response and suppression in the beta band (14-26Hz). Even though we used equivalent stimuli in both modalities, our findings indicate that the underlying neuronal circuitry may not share ubiquitous strategies for stimulus processing.

**Significance statement:** In the visual cortex, gratings can induce robust narrowband gamma oscillations (30-70Hz). These visual stimulus-induced oscillations can further be used as a biomarker for diagnosing neuronal disorders. However, tasks used to elicit these oscillations are challenging for elderly subjects, and therefore, we tested if we could use auditory stimuli instead. We hypothesized that auditory ripple stimuli, which are analogous to visual gratings, may elicit these narrowband oscillations. We found that ripples induce a broadband high-gamma response (70-150Hz) in human EEG, unlike visual gratings that produce robust gamma. Thus, the underlying neural circuitry in the two areas may not be canonical.

## Introduction

Modulations in the gamma band (≥ 30Hz) are associated with higher cognitive processes (Tallon-Baudry et al., 1999; Ray and Maunsell, 2010) and sensory representation (Gray and Singer, 1989; Brosch et al., 2002). Sensory input driven narrowband gamma oscillations (∼30-70Hz) have been identified in various species and neocortical areas (Adrian, 1942; Gray and Singer, 1989; Gieselmann and Thiele, 2008; Muthukumaraswamy and Singh, 2013) and are thought to be due to the reciprocal interaction between excitatory glutamatergic pyramidal neurons and inhibitory GABAergic interneurons (Cardin et al., 2009). Therefore, only the stimuli that drive the neuronal population in a time-synchronized manner can induce such oscillations. Cartesian gratings are considered an archetype to induce sustained narrowband gamma in the visual cortex. The magnitude and peak frequency of these oscillations depend on low-level grating features such as size, contrast, orientation, and spatial and temporal frequency (Gieselmann and Thiele, 2008; Ray and Maunsell, 2010, 2011; Murty et al., 2018). Recent studies have shown that the amplitude of oscillations induced by gratings in electroencephalography (EEG) recording decreases with healthy ageing (Murty et al., 2020) and is weaker in patients with mild cognitive impairment and Alzheimer’s disease (Murty et al., 2021) compared to age and gender-matched healthy control subjects. This suggests that these oscillations can be potentially used as a biomarker for identifying cognitive decline. However, such studies typically require eye fixation and tracking while a full-screen, high luminance-contrast grating is presented on the screen. Therefore, the task becomes challenging as it may lead to visual discomfort (Wilkins et al., 1984), especially for elderly subjects.

Replacing the visual stimulus with an auditory one would resolve such challenges, as subjects can sit with closed eyes and passively listen to auditory stimuli, provided that the auditory stimulus induces a narrowband rhythm. In-vitro studies of rat auditory cortex have been shown to elicit a narrowband oscillation (30-80Hz) in response to stimulation (Ainsworth et al., 2016) and have shown to have distinct generators for lower (30-45Hz) and higher gamma (50-80Hz) (Ainsworth et al., 2011). However, in in-vivo recordings with auditory stimuli such as pure tones (Crone et al., 2001; Brosch et al., 2002; Edwards et al., 2005; Steinschneider et al., 2008; Fujioka et al., 2009), short tone bursts (Trautner et al., 2006; Vianney-Rodrigues et al., 2011), phonemes (Crone et al., 2001; Edwards et al., 2009), clicks (Brugge et al., 2009), frequency sweeps (Jeschke et al., 2008; Lenz et al., 2008), noise (Griffiths et al., 2010; Sedley et al., 2012), words (Canolty et al., 2007), and sentences (Billig et al., 2019), an increase in power occurs mainly at frequencies higher than 60 Hz. These are “broadband” increases and may reach up to ∼200Hz, with unequal power increases across frequencies (Crone et al., 2011). Hence, auditory stimulus-induced narrowband gamma oscillations have not been shown unequivocally.

Even in the visual cortex, narrowband gamma is strongly elicited only by specific stimuli such as bars (Gray and Singer, 1989), gratings (Jia et al., 2011; Murty et al., 2018) and certain iso-luminant hues (Shirhatti and Ray, 2018). Therefore, we tested whether auditory ripple stimuli, whose attributes match that of a visual grating in terms of feature complexity (Shamma, 2001) and neural representation (deCharms et al., 1998), might induce auditory narrowband gamma oscillations (see Discussion for more details). Ripples are generated by superimposing multiple sinusoidally amplitude-modulated tones (Kowalski et al., 1996). They are parametric, meaning they can be fully characterized using limited features that can be changed independently while maintaining their ethological relevance, as their spectra match that of natural vocalizations (Langers et al., 2003). We recorded 64-channel EEG from human subjects who passively listened to ripple sounds from a loudspeaker with their eyes open or closed or passively fixated on the computer screen during the presentation of a full-screen grating stimulus on the monitor and compared the gamma responses generated by these stimuli.

## Methods

### Human Subjects

We recruited 36 healthy subjects (aged 22-38 years, a mean of 26.6±3.7 years, 18 females) for the study from the Indian Institute of Science community. Participants reported having normal hearing levels with no abnormalities and had corrected to normal vision (except for one participant with strabismus). All subjects, barring one, were right-handed. Participation was voluntary, and written informed consent was obtained from all the subjects after briefing them about the experimental procedure. Subjects were given monetary compensation for their time and effort. Experiments were performed according to the protocol approved by the Institutional Human Ethics Committee of the Indian Institute of Science, Bangalore.

### EEG setup and data acquisition

Raw EEG signals were recorded from 64-channel active electrodes (actiCap) using the BrainAmp DC EEG acquisition system (Brain Products GmbH). Electrodes were placed using the international 10-10 standard reference scheme with FCz as the reference electrode (unipolar reference scheme). Raw signals were sampled at 1000Hz and were filtered between 0.016Hz (first-order filter) and 250Hz (fifth-order Butterworth filter). Signals were digitalized at 16-bit resolution (0.1µV/bit). Impedance values were kept below 25kΩ for the entire recording duration. Average impedances of the final set of electrodes were 6.88±3.06 KΩ (mean± std) for the visual task, 6.76±2.99 for the eye-close auditory task and 7.29±3.02 KΩ for the eye-open auditory task.

### Experimental setting and task

All thirty-six subjects did the visual and eye-close auditory tasks; twelve subjects out of these also participated in the eye-open auditory task. The first twenty-four (twelve females) subjects performed the visual protocol first, followed by the auditory protocols, which comprised the eye-close version and two other auditory protocols (not described in this study). For these participants, auditory protocols were run in a counterbalanced order. The remaining twelve subjects completed the visual, eye-close and eye-open auditory task versions. These protocols were counterbalanced.

#### Visual Task

Each participant performed a passive fixation task. They sat in a dark room in front of a gamma-corrected LCD monitor (BenQ XL2411; resolution 1280A×720 pixels; refresh rate 100Hz; mean luminance 60 cd/m^2^). The monitor was placed 58cm away from the subject’s eyes, so the full-screen stimulus subtended the width and height of 49.4° and 29° of the visual field. The visual stimuli, sinusoidal luminance grating, were presented using the NIMH MonkeyLogic software tool on MATLAB (The MathWorks, Inc; <u>RRID: SCR_001622</u>). The marker for stimulus onset was recorded in the EEG file by using a digital I/O card (National Instruments USB 6008 or USB 6210 multifunctional I/O device).

Since the auditory task (see below) involved the presentation of a continuous sequence of auditory stimuli with some inter-stimulus interval, we modified our visual task to be comparable to the auditory task. Specifically, unlike our previous studies (Murty et al., 2018, 2020, 2021), we presented visual stimuli in a single long continuous sequence where each stimulus was presented for 800ms followed by an inter-stimulus interval of 700ms. A square fixation spot of 0.2° was shown in the center of the screen throughout the task duration. Subjects were instructed to hold and maintain their fixation whenever the visual stimulus was presented and blink or break fixation (if needed) during the inter-stimulus interval. Eye position was monitored continuously (see details below), and epochs with breaks in fixation were discarded offline later. Visual stimuli were achromatic luminance full contrast and full-screen gratings presented at one of two spatial frequencies (SF - 2 and 4 cycles per degree) and one of four orientations (Ori - 0°, 45°, 90° and 135°), generating eight different kinds of stimuli. Each stimulus type was repeated 25 times, yielding 200 trials. We chose these stimulus parameters as they induce a robust gamma response (Murty et al., 2018). The total duration was ∼5 minutes. For a couple of subjects, the stimulus sequence was paused for 30-60 seconds because they requested a break, after which the sequence was resumed.

#### Auditory Task

The task had an eye-close and eye-open version. In the eye-close session, subjects sat in a dark room with closed eyes and were instructed to listen passively to the sounds. They were instructed to keep their eyes closed to minimize eye movement or blink artefacts. To make the auditory task equivalent to the visual task, we ran an eye-open version on a subset of subjects, where subjects had to fixate on the screen where a fixation spot of 0.2° was shown at the center, and sounds were played in the background. The sessions were conducted in a quiet room to minimize irrelevant sounds. The stimuli were played using a multidirectional speaker (Marshall Kilburn II) at 75-80 dB. The sound stimuli were generated in MATLAB by custom-written code. They were presented using the NIMH MonkeyLogic software tool on MATLAB. The marker for stimulus onset was recorded in the EEG file by using a digital I/O card (National Instruments USB 6210 multifunctional I/O device).

Each trial had one stimulus of 800ms followed by an inter-trial interval of 800ms. The stimulus set consisted of spectro-temporally modulated ripples. Ripple stimuli have a broadband carrier with a sinusoidally varying spectral envelope that drifts along the logarithmic frequency axis at a constant velocity (Kowalski et al., 1996). The stimuli composed of 80 tones equally spaced along the logarithmic axis, spanning 5 octaves (250-8000Hz, 16 tones per octave), sampled at 44100Hz, were used. The amplitude of all individual tones was sinusoidally modulated on a linear scale and was modulated at 90% or 10dB. Each ripple stimulus had 10ms – on/off ramps. The stationary ripple stimuli were presented at one of three spectral modulation frequencies or ripple density (Ω - 0, 0.8 and 1.6 cycles/octave). A stationary ripple can be represented mathematically as –

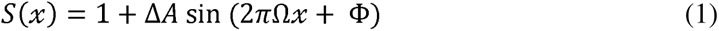

where ΔA, modulation depth, is 0.9; x is the logarithmic frequency axis (in octaves), defined as 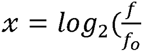 where *f*_0_ is the lower frequency edge, i.e., 250Hz and *f* is the component tone frequency. The starting phase of the ripple (Φ) was defined relative to the low low-frequency edge of the spectrum and kept at 0. The moving ripples were obtained by temporally modulating stationary ripples, such that the envelope moved downwards towards lower frequencies at one of the four velocities constantly (*ω* - 0, 5, 10, 20 cycles/second (Hz)). The moving ripple spectro-temporal spectrum can be represented by –

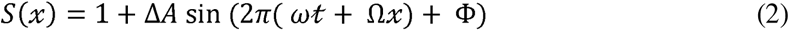

We chose these stimulus parameters based on the previous study done with ripple stimuli in humans (Langers et al., 2003). This generated 12 possible combinations, and each stimulus type was repeated 40 times, totalling 480 trials. The total task duration was about ∼15 minutes, divided into two blocks with 2–3-minute breaks in between according to the subject’s comfort.

Owing to their spectro-temporal response profile, ripple envelopes look like visual gratings and are thus often described as acoustic analogues of visual gratings (Shamma, 2001). The spectral modulation frequency can be considered equivalent to spatial frequency, which determines how dense the gratings are. The modulation depth of the ripple envelope is analogous to the contrast of visual gratings; similarly, temporal modulation is like the temporal frequency of drifting visual gratings. Given the similarity between features, ripples are referred to as auditory gratings. In the rest of the paper, we will refer to these stimuli as auditory gratings. Our rationale for using auditory gratings is discussed in more detail in the Discussion.

### Eye position Analysis

Eyes signals were recorded either using Eyelink Portable Duo head-free eye tracker or Eyelink 1000 (SR Research Ltd, sampled at 1000Hz) for the entire duration of the visual and eye-open version of the auditory task. Before the start of each session, the eye tracker was calibrated for pupil position and monitor distance. No online artefact rejection was done. During analysis, we segmented epochs of EEG data between -848ms to 1650ms around each stimulus onset. We obtained 200 and 480 such segments for the visual and auditory tasks, respectively. Each segment is referred to as a “trial”, even though this task had no trial structure as the stimuli were presented continuously. We rejected trials offline, which had fixation breaks, defined as eye blinks or shifts in eye position outside a square window of width 5° centred on the fixation spot during -500ms to 750ms of stimulus onset. This interval was chosen to ensure that the “baseline” (-500ms to 0ms of stimulus onset) and “stimulus” (250ms to 750ms) periods used for the calculation of the power spectral density (PSD) were both free of eye-movement-related artefacts. It led to the rejection of 20.58±16.72% (mean ± std) trials from the visual task, barring one subject for which all trials were labelled bad as the subject had strabismus. For the auditory eye-open task, we rejected 19.46±16 % of trials.

### Artefact Rejection

We used a fully automated artefact rejection pipeline (see (Murty and Ray, 2022), for details) with one minor modification. In our previous method, the threshold for rejection was based on the standard deviation (SD) in the time-series data (any trial in which any time point between -0.5 to 0.75 seconds deviated by more than 6 SD was rejected), but this threshold could vary depending on the outliers. Here, we chose hard cutoffs, which helped us evade this issue altogether. Specifically, after rejecting all electrodes with impedance >L25kΩ and bad eye trials (as done in the previous study), we first calculated the root mean squared (RMS) value for each trial (-0.5s to 1.5s) after passing it through a high pass filter of 1.6Hz (to remove any slow drifts), and applied a fixed threshold bound (upper RMS cutoff of 35µV and lower RMS cutoff of 1.5µV) and labelled any trial that lay outside that bound as bad for that electrode.

Next, we computed multi-tapered PSD for the rest of the trials (using the Chronux toolbox; (version 2.10) (Bokil et al., 2010), <u>RRID: SCR_005547</u>), available at http://chronux.org). Any repeat for which the PSD deviated by six times the standard deviation from the mean at any frequency point (between 0 -200Hz) was also labelled bad. After this, we listed a common set of bad repeats across all 64 electrodes. We discarded the electrodes with more than 30% of all repeats labelled as bad. Any trial was labelled bad if it occurred in more than 10% of the remaining electrodes. Next, we selected a subset of occipital, parieto-occipital (O1, Oz, O2, P1, P2, P3, P4, PO3, POz, PO4 – set used in Murty et al., 2018, 2020) and temporal electrodes (TP9, TP7, TP8, TP10), and any trial that was marked as bad in any of these electrodes were also included in the set of common bad trials. This was done to ensure that the electrodes used for the calculation of power for the visual and auditory conditions were artefact-free.

These criteria led to the rejection of less than ∼30% (28.5± 9.1%) of the data collected for the eye-close auditory task. For the visual and eye-open auditory tasks, these criteria and offline eye rejection led to the rejection of 36.73 ± 12.89% and 41.90 ± 11.06% of data, respectively. Note that this relatively higher percentage of bad trials compared to our previous studies is due to the continuous stimulus presentation paradigm used in this study. In previous studies, a trial had 2-3 stimuli, after which the subjects could break fixation or blink their eyes. But here, the stimulus presentation was in a continuous stream, and subjects were allowed to blink (if needed) during the inter-stimulus interval; hence, more data segments (“trials”) had to be discarded.

As an additional criterion to reject electrodes, we calculated the slope in the range of 56-86Hz of baseline PSD (averaged across all good trials) for each electrode by fitting the PSD with a power-law function (for details, refer to Murty et al., 2020). We discarded any electrode for which the PSD (1 taper) slope was less than 0. Even after applying stringent conditions to reject electrodes, if some electrode that had not been rejected yet was visually noisy, we manually declared that electrode as bad. We removed an additional 1.7% of the total electrodes by doing this.

We then rejected any subject from the analysis of visual or auditory tasks with less than 50% of the occipital (O1, Oz, O2, PO7, PO3, POz, PO4, PO8, Iz) or temporal (FT9, FT7, T7, TP7, TP9, FT10, FT8, T8, TP8, TP10) group electrodes, respectively. This way, we rejected two participants from the eye-close auditory task and one from the eye-open auditory task. For the visual task, 2 participants were rejected as one had strabismus, and the baseline signal was extremely noisy for the other.

### EEG data analysis

As in our previous study (Murty et al., 2020), we used both unipolar and bipolar reference schemes for analysis. For the unipolar reference scheme, we considered the following electrodes – TP9, TP7, TP8, and TP10 for auditory task analysis and P1, P2, P3, P4, PO3, POz, PO4, O1, Oz, O2 for visual task analysis. For the bipolar referencing scheme, we re-referenced data from each electrode data to its neighbouring electrode. We thus obtained 114 bipolar pairs out of 64 unipolar electrodes. We have used the following bipolar combinations – TP9–TP7, TP7–T7, TP7–P7, TP7–CP5, TP10–TP8, TP8–T8, TP7–P8, TP7–CP6 for auditory task analysis and PO3–P1, PO3–P3, POz-PO3, PO4–P2, PO4–P4, POz-PO4, Oz-POz, Oz-O1 and Oz-O2 for visual task analysis. Depending upon the reference scheme, data were pooled for all these electrode or electrode combinations.

All the data were analyzed using custom-written codes in MATLAB. Using the Chronux toolbox, we computed multi-tapered PSD and time-frequency spectrograms using a single taper. We chose a period between -500ms and 0 ms (0ms marks the stimulus onset) as the baseline and a period between 250ms to 750ms as the stimulus period, with a frequency resolution of 2Hz. For spectrograms, we used a moving window of size 250ms and a step size of 25ms, thus yielding a frequency resolution of 4Hz. We calculated the change in power for narrowband gamma oscillation (*f* E l 20 - 66J Hz) and high-gamma activity (*f* E l 70 -150JHz) as follows:

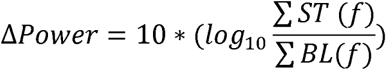

*ST* and *BL* represent power across frequency(*f*) for stimulus and frequency averaged across all stimuli repeats for a particular condition and electrodes.

Scalp maps were generated using the topoplot.m function of the EEGLAB toolbox ((Delorme and Makeig, 2004) <u>RRID: SCR_007292</u>). The function was modified to show each electrode as a coloured disc, with the colour representation change in power for a particular frequency range in decibels (dB).

### Statistical Analysis

We compared the means of the subject averaged power spectral density during stimulus and baseline periods using a paired t-test test. For comparing the change in power across visual and auditory protocols, unpaired t-tests with unequal variance and F-statistic were used.

### Data and Code availability

Spectral analyses of the data were performed using the Chronux toolbox. Raw data will be made available to readers upon reasonable request.

## Results

We collected 64-channel EEG data from 36 subjects (18 females). Participants passively listened to auditory grating stimuli or passively fixated on the screen while presenting full-screen sinusoidal grating stimuli on a monitor. Each stimulus was 800ms long in either modality, preceded by a baseline period of 800ms for auditory and 700ms for visual stimuli.

Fig.1 shows the subject averaged time-frequency spectrograms for change in power from baseline (-500 to 0ms of stimulus onset) for each electrode as positioned on the scalp for visual (right) and auditory protocol (left). The spectrograms are averaged across all stimulus conditions. Visual grating stimuli elicited a robust narrowband gamma response in occipital electrodes (in the range of 22-64Hz) and a pronounced alpha (8-14Hz) suppression. However, auditory gratings did not produce such a narrowband gamma response. In contrast, they induced a prominent high-gamma activity (70-150Hz) in electrodes located near mastoids and suppression in beta rhythms (14-26Hz) across all electrodes. The response elicited by the auditory gratings was much weaker than that elicited by the visual grating stimuli (note the difference in scale in A versus B). For visual stimuli, some frontal electrodes showed a broadband response after stimulus offset, likely due to artefacts related to eye movements or blinks (we removed only those trials for which eye movement occurred between -500 to 750ms of stimulus onset; see Methods for more details). This frontal response was also observed for the auditory protocol when eyes were open, as shown later.

**Figure 1:**
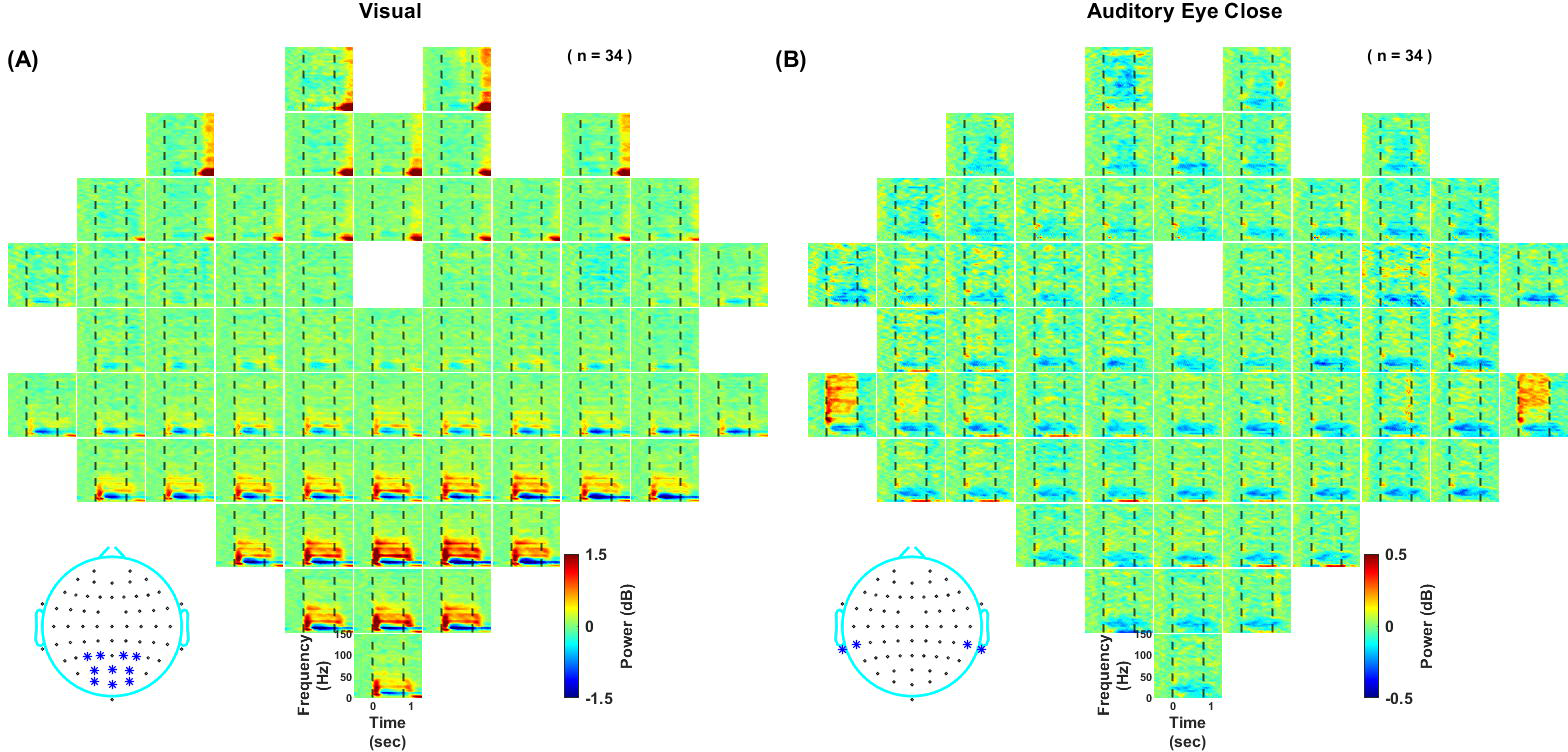
Change in power after stimulus presentation pooled across all stimulus conditions. Subject-averaged time-frequency change in power spectra up to 150Hz for all 64 channels (frequency: 0-150Hz, vertical axis and time: -500-1300ms, horizontal axis). The plots are arranged according to the 64-channel layout (actiCap) with FCz as the reference electrode, unipolar referencing. Stimulus is presented during 0 to 800ms, indicated by dashed vertical lines. Colorbar indicates log Power ratio in decibels (dB). **(A)** In response to the presentation of the visual grating stimuli. **(B)** In response to the presentation of the auditory grating stimuli. The star-marked electrodes in the topoplots at the bottom show the electrodes used for further analysis.

Next, we averaged the responses across selected electrodes (as shown in Figure 1 inset; see EEG data analysis section in Methods) depending upon the stimulus modality. Fig 2A (Top row, left panel) shows that visual grating stimuli elicit two distinct gamma bands, termed slow gamma (∼20-34Hz) and fast gamma (∼36-66Hz) in spectrograms (Murty et al., 2018, 2020). On the other hand, auditory grating stimuli elicited a high-gamma response (70-150Hz), Fig 2A (Bottom row, left panel). Our previous studies showed that using a bipolar referencing scheme improved the narrowband gamma response (Murty et al., 2020), so we performed the same analysis using bipolar referencing as well (Figure 2A; right panel). There was a marginal improvement in the visual-induced narrowband gamma response, especially in the fast gamma band. However, the auditory stimulus-induced high-gamma activity weakened with bipolar referencing (Fig. 2A bottom panels). The change in power in the narrowband gamma range was about ∼1dB when visual stimuli were presented (Figure 2B, second row), which was missing for auditory stimuli (Figure 2B, fourth row). Conversely, the change in high-gamma power was ∼0.05 dB for visual stimuli and ∼0.1dB for auditory stimuli for the unipolar case (note the difference in scales in the plots). The topoplots in Fig 2C showed that visually induced gamma was localized in the occipital and parieto-occipital region, while auditory stimuli induced broadband high-gamma in electrodes located near the mastoids. For further analysis, we have used unipolar referencing.

**Figure 2:**
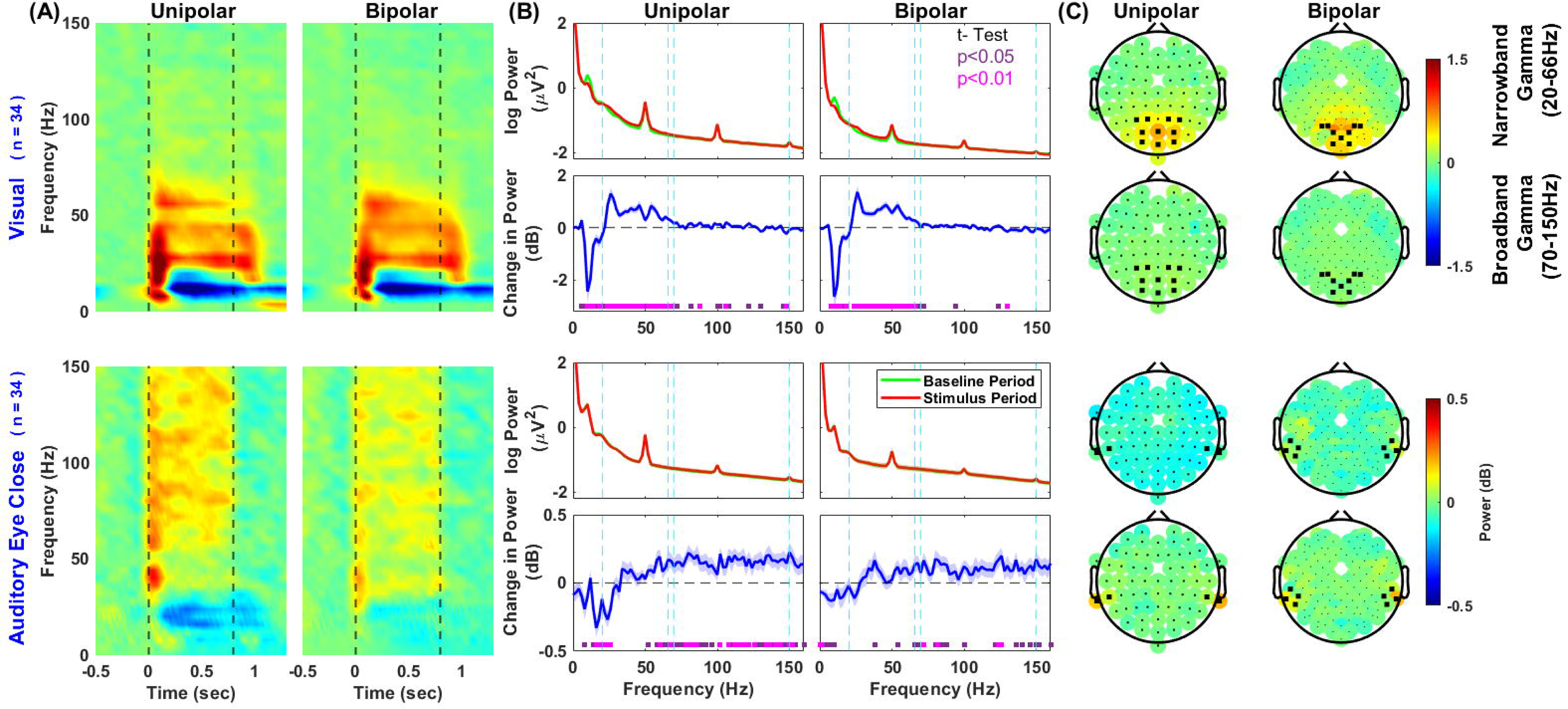
Subject and electrode averaged change in power compared to baseline (- 500ms to 0ms), pooled across all stimulus conditions with two different reference schemes. Results are shown for the unipolar reference scheme (left) and bipolar (right). The top row in A and the first two in B and C are in response to visual stimuli. The bottom row in A and the last two in B and C are in response to the auditory stimuli. Electrodes used for averaging in A and B are highlighted with black dots in scalp maps shown in C. **(A)** Time-frequency change in power. Dashed vertical lines (black) represent stimulus onset and offset. **(B)** Power spectra (first and third-row panels – red trace is stimulus period (250ms to 750ms), and the green trace is the baseline period (-500ms to 0ms)) and change in power spectra vs frequency (second and fourth-row panels, blue traces). The solid traces represent the mean across subjects, and thickness represents SEM. Dashed vertical lines represent narrowband gamma (20-66Hz, blue) and broadband high-gamma (70-150Hz, cyan). Coloured squares at the bottom represent the significance of differences in means (paired t-test – purple: p-values between 0.01-0.05, pink: p <0.01). **(C)** Scalp maps of 64 unipolar electrodes (left) and 114 bipolar electrodes (right). The first and third rows show the change in power for frequencies 20-66Hz. The second and fourth rows show the change in power for frequencies 70-150Hz. The colorbar represents the log power ratio in dB.

### Gamma and high-gamma responses were uncorrelated across subjects

Figure 3 shows the change in power spectrograms (for the same set of visual and auditory electrodes as Figure 2) for individual subjects, sorted by decreasing auditory high-gamma power. Although auditory high-gamma was much weaker than visual narrowband gamma, it appeared stronger than visual high-gamma. The range of auditory stimuli induced broadband high-gamma activity varied among the subjects. Further, visual and auditory stimuli had no consistent effects on subjects – subjects with strong auditory high-gamma did not have stronger visual narrow or broadband high-gamma, or vice-versa.

**Figure 3:**
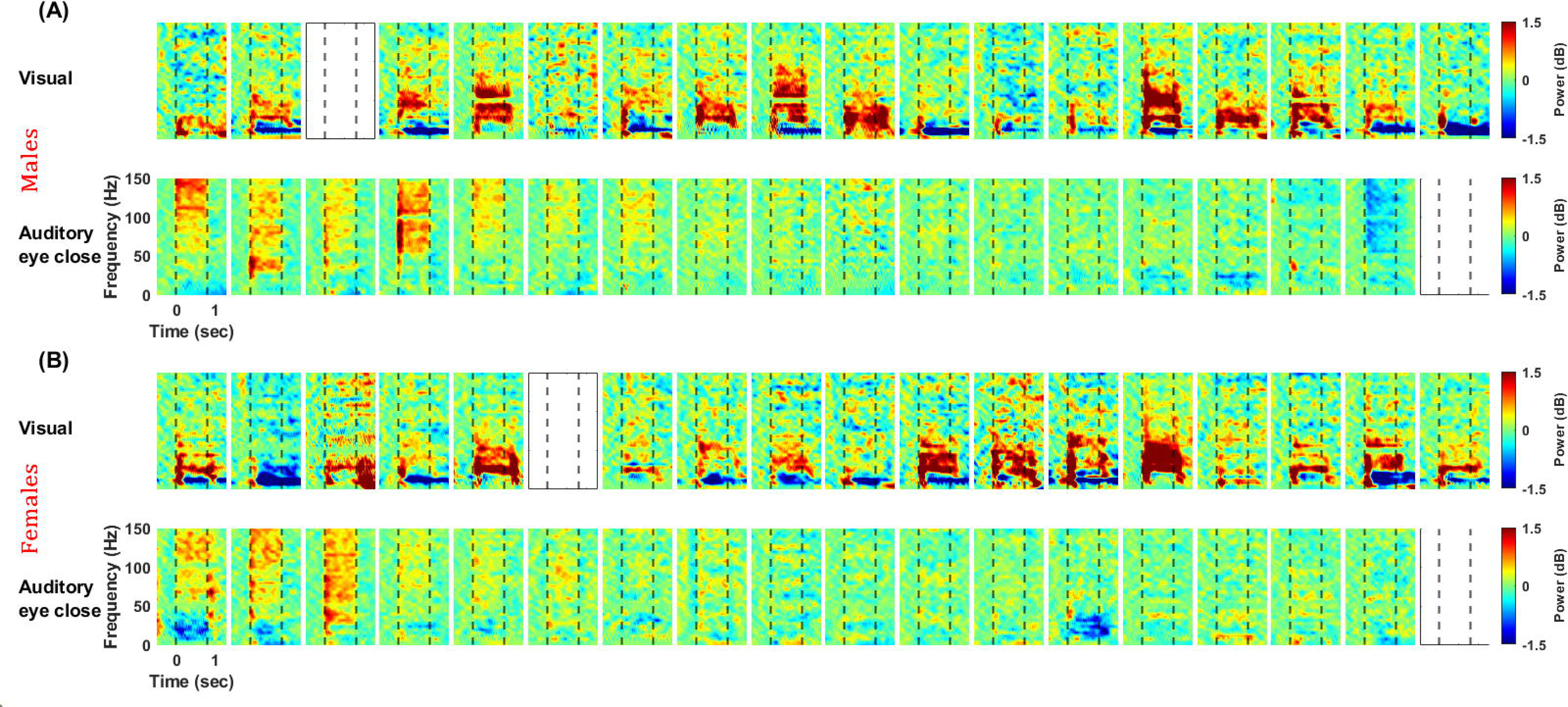
Visual and auditory responses for individual subjects. Change in time-frequency power spectra for males **(A)** and females **(B)**. The top row represents the results in response to visual stimuli, and the bottom represents the results in response to auditory stimuli. Subjects are ordered horizontally based on the decreasing high-gamma activity in 70-150Hz for the auditory condition, starting from the left. The response is averaged across selected occipital and temporal electrodes (highlighted in the inset of Fig 1) for respective protocols. The colorbar represents the log power ratio in dB. Empty plots refer to subjects rejected from the analysis (refer to methods – artefact rejection for more details).

Figure 4 shows that visual narrowband gamma was significantly stronger than auditory narrowband gamma, which was negligible (Visual: 0.4±0.09, Auditory: -0.06±0.02 (mean±sem), p = 8.6×10^-5^, N = 34, unpaired t-test, F = 15.33). In contrast, broadband high-gamma showed the opposite trend, but the difference was not significant (Visual: 0.061±0.02, Auditory: 0.12±0.03 (mean±sem), p = 0.118, N = 34, unpaired t-test, F = 0.3646, Fig 4E). Since the auditory response was computed over fewer electrodes, the visual high-gamma might have been high for a subset of selected visual electrodes, but the mean value could have been reduced when averaged over all the selected electrodes. To rule this out, we performed the same analysis after taking only a single “best” electrode for each subject with the most robust response for each modality in the respective frequency range (Fig 4C and 4D). The trends remained similar, with the broadband auditory response now significantly higher than visual (narrowband gamma:Visual: 0.80±0.13, Auditory: 0.05±0.03 (mean±sem), p = 1.4×10^-6^, N = 34, unpaired t-test, F = 16.65; broadband high-gamma: Visual: 0.21±0.02, Auditory: 0.36±0.06 (mean±sem), p = 0.0189, N = 34, unpaired t-test, F = 0.1477, Fig 4E). Further, Pearson’s correlation between the responses to visual and auditory stimuli was insignificant for any condition (p-values are shown in the plots in 4A-D).

**Figure 4:**
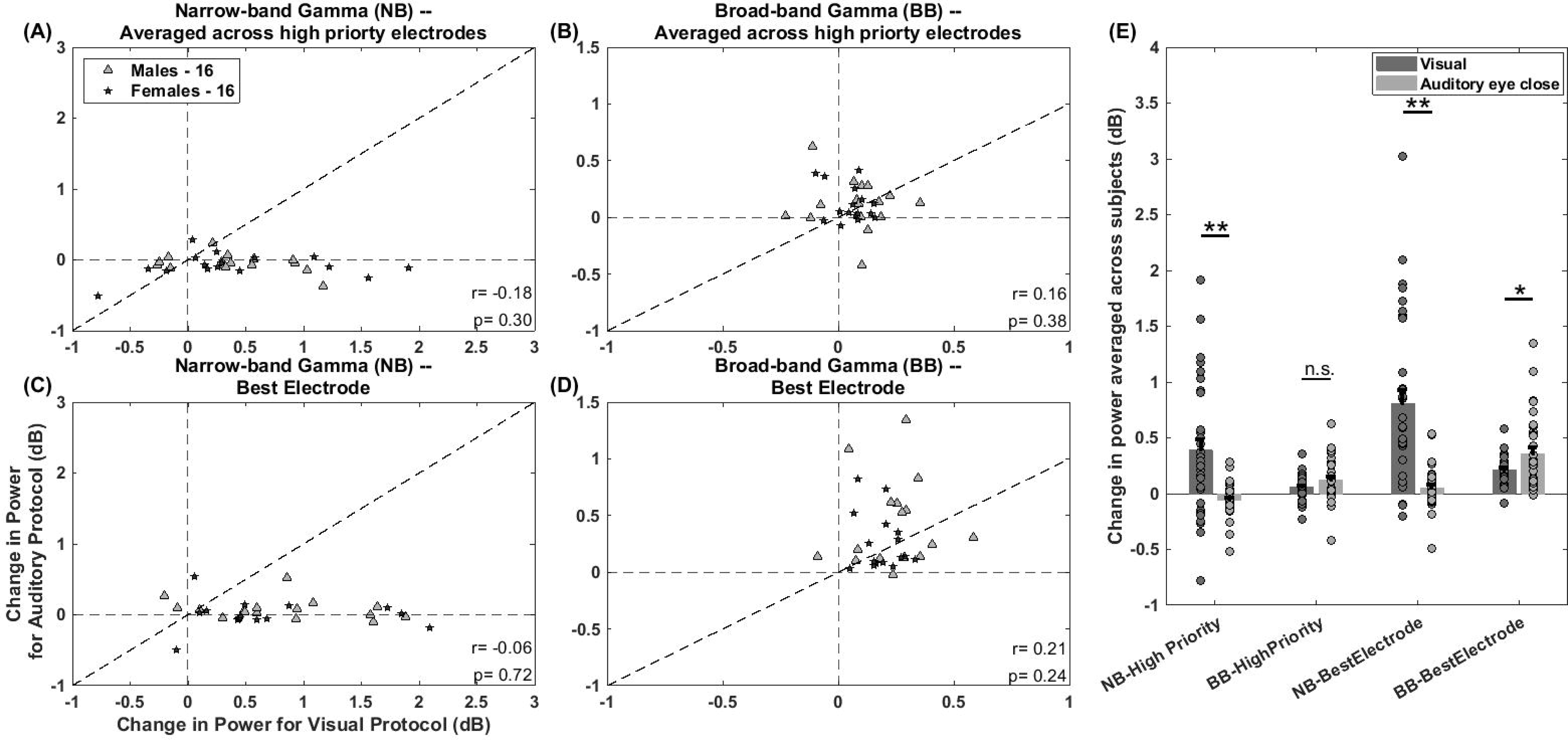
Auditory-induced high-gamma activity is not correlated with visually induced narrowband gamma. Scatter plots for change in power in response to visual stimuli vs auditory stimuli. **(A)** For narrowband gamma power averaged across chosen electrode groups. **(B)** For broadband high-gamma power averaged across chosen electrode groups. **(C)** For narrowband gamma power for electrode with maximum change in power. **(D)** For broadband high-gamma power for electrode with maximum change in power. Pearson’s correlation coefficient and p-value are mentioned at the bottom right in each panel. Note the difference in axis limits across the panels. **(E)** Bar plots showing the mean change in power from baseline across subjects for different gamma frequency ranges (NB refers to Narrowband gamma and BB refers to Broadband high-gamma) during different tasks. The dots represent the change in the power of each subject. The significance of the unpaired t-test (with unequal variance) is indicated at the top of the bar plots.

### No tuning characteristics or stimulus selectivity could be observed in EEG

The previous results were the average responses across eight and twelve different stimuli for visual and auditory conditions, respectively (see Methods for details). To test whether gamma/high-gamma was selective for some stimuli, we plotted the change in time-frequency power spectra for different stimulus conditions (Figure 5). We did not observe gamma activity tuned to any particular stimulus in either of the modalities. To eliminate the possibility that individual subjects were tuned to different stimulus features, we determined their coefficient of variation (CV) of power across stimulus conditions. Specifically, during the stimulus period (250ms to 750ms), we calculated a ratio of average power in narrowband gamma and broadband high-gamma frequency ranges across stimulus conditions to its standard deviation for each selected electrode, depending on the task for each subject. The values were then averaged across electrodes. The values were generally small (Narrowband: Visual: 0.09±0.008, Auditory: 0.071±0.005 (mean±sem); p = 0.0475, N=34, unpaired t-test, F = 2.12; Broadband: Visual: 0.05±0.003, Auditory: 0.066±0.004 (mean±sem); p = 0.0475, N=34, unpaired t-test, F = 0.6047) indicating poor selectivity towards any stimulus feature. These results are consistent with our previous study where we found strong orientation selectivity for narrowband gamma in local field potential (LFP) data from monkeys, which was absent in human and monkey EEG (see Figure 2E of Murty et al., 2018).

**Figure 5:**
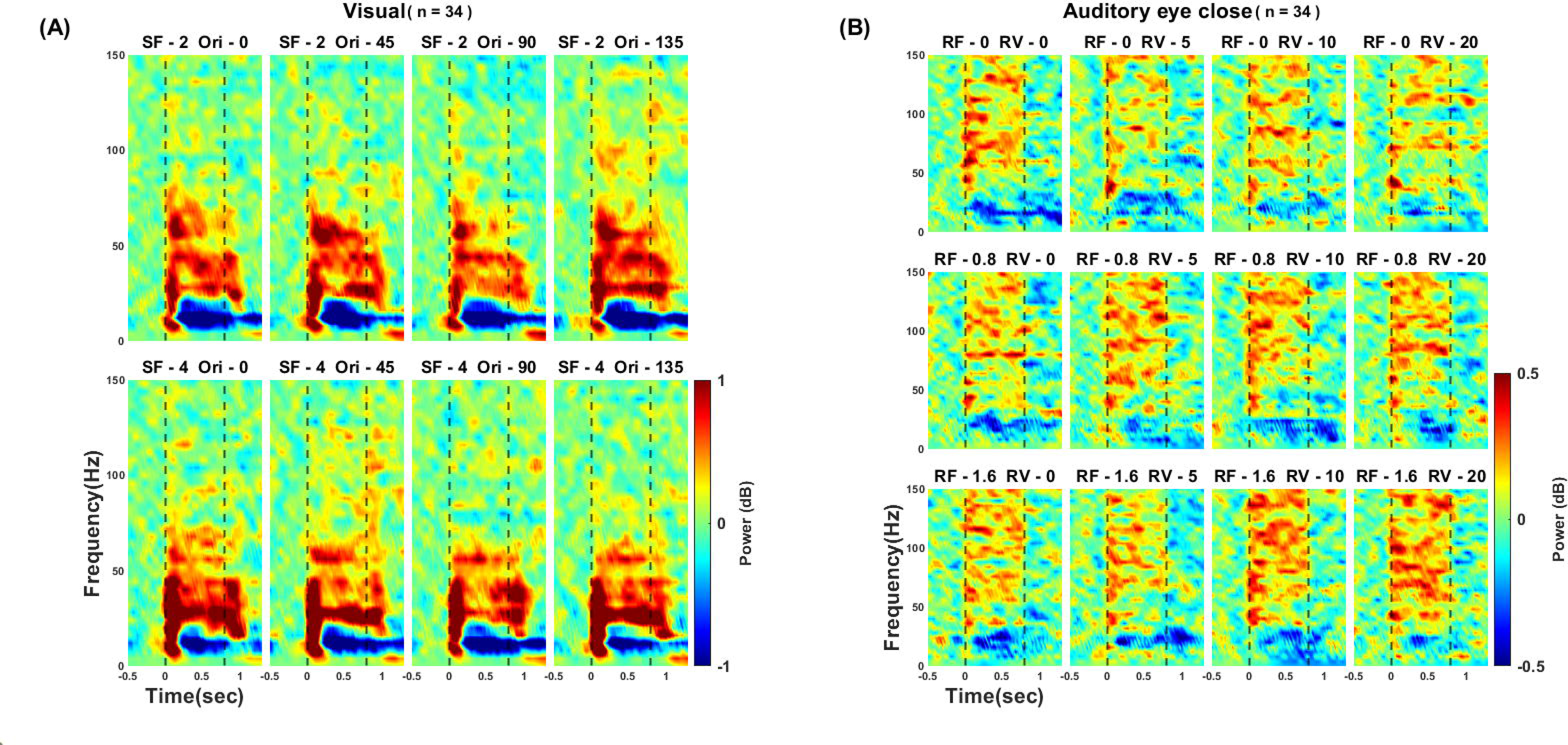
No stimulus selectivity was observed for either of the induced gamma(s). Change in time-frequency power spectrum from baseline for different stimulus conditions. The last row and column represent averaged responses across that row or column. **(A)** In visual modality. **(B)** In auditory modality. Power is averaged across chosen electrodes for each modality. Dashed vertical lines (black) represent stimulus onset and offset. The colorbar represents the log power ratio in dB.

### Broadband High-gamma activity remains the same for the eye-close and eye-open auditory tasks

Sensory-driven gamma oscillations are known to be modulated by arousal (Vinck et al., 2015). Since subjects had their eyes closed for the auditory task, whereas they were instructed to passively fixate on the monitor for the visual task, we ran an eye-open version of the auditory task where subjects passively fixated on the screen while passively listening to the sounds to ensure a similar arousal level. We found a similar increase in high-gamma activity and suppression in the beta band for the auditory eye-open task as well (Figure 6). We also saw increased broadband activity after stimulus offset attributed to eye artefacts, similar to those observed in the visual task. The high-gamma power was comparable for eyes-open and eyes-closed cases: (for high-priority electrodes: Eyes-open: 0.093±0.05 and Eyes-closed: 0.116±0.11 (mean±sem), p = 0.7145, N = 11, unpaired t-test, F = 1.93; for best electrode: Eyes-open: 0.297±0.03 and Eyes-closed: 0.346±0.11 (mean±sem), p = 0.7592, N = 11, unpaired t-test, F = 0.88). The power values were significantly correlated for the best electrode, for which maximum change in power was calculated. (Pearson’s linear coefficient: r = 0.75, p = 0.0079).

**Figure 6:**
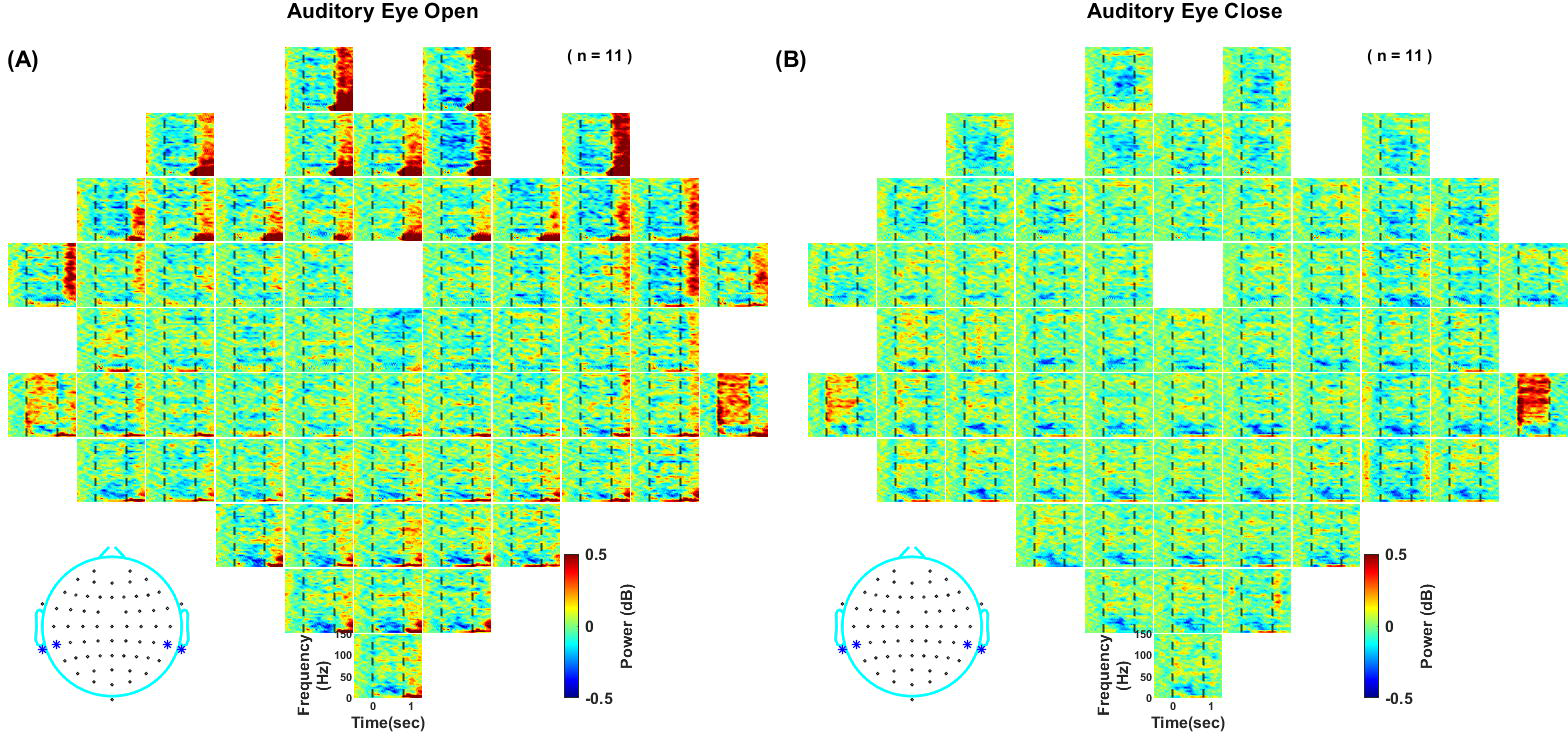
**Subject-averaged time-frequency change in power spectra** for the **(A)** eye-open and **(B)** eye-close auditory tasks—same format as Figure 1.

## Discussion

We tested whether narrowband gamma oscillations can be induced by auditory grating stimuli analogous to that induced by visual sinusoidal luminance gratings in the visual areas in human EEG during passive stimulus presentation tasks. We found that the auditory gratings induced an extremely focal broadband high-gamma response (∼70-150 Hz) in temporal electrodes and a widespread decrease in beta rhythm (∼14-26Hz). In contrast, visual grating stimuli induced narrowband gamma oscillations (∼20-70Hz) in the occipital and parietooccipital electrodes accompanied by suppression in alpha oscillations (∼8-14Hz) in the same subjects. The auditory grating-induced broadband activity was weaker than narrowband oscillations elicited by the visual gratings but was still robust than their broadband response. Subjects which showed an induced response to either stimulus did not respond similarly to the other stimulus modality, indicating that the networks responsible for the induced responses work independently in each modality. We observed poor tuning of these responses towards specific stimuli of either modality, consistent with poor stimulus selectivity for visual gamma observed earlier in human EEG (Murty et al., 2018).

### Comparison with previous studies

Several studies have reported auditory broadband high-gamma activity in response to a multitude of stimuli and various aspects of auditory cortical processing in primates. When simple stimuli such as pure tones (Crone et al., 2001; Edwards et al., 2005; Steinschneider et al., 2008; Fujioka et al., 2009), tone bursts (Trautner et al., 2006), and clicks (Brugge et al., 2009) were used, only a transient increase in broadband high-gamma was reported. However, our study obtained sustained high-gamma responses throughout the stimulus presentation; this might have been because the neurons in the auditory cortex respond better to complex stimuli than pure tones (Tian and Rauschecker, 2004) and are capable of firing in a sustained fashion only when optimal stimuli are used (Wang et al., 2005). Thus, auditory gratings are better suited to drive neurons in auditory areas. These results also align with the results of other studies which used complex stimuli, such as frequency sweeps (Lenz et al., 2008), phonemes (Edwards et al., 2009), words (Canolty et al., 2007), and sentences (Billig et al., 2019). A recent study (Ross et al., 2020) also reported increased evoked high-gamma activity (90-150Hz) in response to a syllable presentation in human EEG. The polarity of response at electrodes TP9 and TP10 electrodes was opposite to the results that we obtained. However, these differences could be accounted for by the fact that the location of the reference electrode affects EEG measurements (Nunez and Srinivasan, 2006); in our study, it was located at FCz and was located on the nose in their study.

Narrowband gamma oscillations to auditory stimuli have only been reported in studies performed on rodents. In-vitro studies done on rat A1 neocortical slices have shown the presence of narrowband gamma in the range of 30-80 Hz in isolation (Ainsworth et al., 2016), i.e. without any broadband activity and further showed that two distinct local networks can give rise to such oscillations in this frequency range (Ainsworth et al., 2011). However, studies in awake rats and Mongolian gerbils have shown narrowband gamma response (∼30-70 Hz) along a broadband increase in high-gamma activity till 150 Hz (Jeschke et al., 2008; Vianney-Rodrigues et al., 2011). Lenz and colleagues (2008) repeated the study conducted in gerbils using the same task and stimulus set in human EEG, and they observed an increase in frequencies from 100Hz up to 250Hz without any narrowband gamma oscillation at lower frequencies. The authors indicated that this may reflect species-specific differences and that the underlying neuronal generators may differ. So, we must be careful while assuming the generalizability of results across species.

The studies that involved the presentation of speech stimuli (Canolty et al., 2007; Edwards et al., 2009; Billig et al., 2019) also reported beta desynchronization, similar to the decrease that we observed. We also note that beta desynchronization was observed when participants performed target detection tasks with simpler auditory stimuli, which was a widespread signal across many electrodes in the EEG (Mazaheri and Picton, 2005), similar to our results. Given that the spectrotemporal envelope of auditory gratings is similar to that of speech and can mimic formant transitions between the vowels and accommodate pitch changes (Langers et al., 2003), a decrease in beta might point towards the activation of circuits involved in complex stimuli and/or task processing.

### Mechanisms of gamma oscillations and high-gamma activity

Periodic activation of neuronal assemblies at an optimum time window gives rise to gamma cycles (Buzsáki and Wang, 2012). Previous studies have pointed out a key role of interneurons in the generation of narrowband gamma oscillations in the sensory cortex (Whittington et al., 1995). Synaptic inhibition comes from fast-spiking parvalbumin-expressing and regular-spiking somatostatin-expressing GABAergic interneurons, which generate gamma oscillations in different frequency ranges (Chen et al., 2017). On the other hand, high-gamma activity is thought to reflect population firing near the microelectrode for invasive recordings and synchronous firing for macrosignals such as electrocorticogram (Ray et al., 2008). It, therefore, has distinct origins compared to narrowband gamma (Ray and Maunsell, 2011). Broadband responses are a robust indicator of neuronal firing in the auditory cortex as well (Manning et al., 2009).

### Auditory grating as an effective stimulus to produce narrowband gamma

As discussed in the Methods, auditory gratings have several properties that make them analogous to visual gratings. In addition, we thought that they would be good candidates for generating narrowband gamma for two reasons. First, they drive the auditory neurons very strongly (deCharms et al., 1998). Visual stimuli that generate narrowband gamma in the visual cortex, such as bars and gratings, also drive the neurons in the visual cortex strongly, and in fact, the ideal filters for primary visual cortex (V1) neurons have oriented Gabor-like features (Olshausen and Field, 1996). It is possible that when the local orientation of the grating of a particular spatial frequency matches the preference of the local cortical neurons, the strong excitatory drive can induce narrowband gamma. Similarly, early processing of sounds and images share equivalent filter characteristics, and the sensory code in the auditory system prefers broadband sounds with smooth edges (Lewicki, 2002). The tuning of the primary auditory cortex for spectro-temporal frequencies of auditory gratings matches the tuning found for spatial frequencies in the primary visual cortex (Kowalski et al., 1996).

Second, narrowband gamma in the visual cortex becomes stronger when the stimulus size increases because of local inhibition generated by surround suppression (Gieselmann and Thiele, 2008). In the auditory cortex, frequency information is spatially coded in tonotopic maps; therefore, we speculated that broadband auditory gratings, which have power over five octaves (250 – 8000Hz), to excite a large neuronal population in the auditory cortex, replicating the effect of large-sized visual grating. Furthermore, auditory cortical neurons can lock to the amplitude modulations of the envelope of the auditory gratings and thus produce sustained responses (Elhilali et al., 2004). Further, the inhibitory circuitry in visual and auditory cortices has similar properties. For example, PV^+^ interneurons in visual and auditory areas are narrowly tuned for spatial frequency and orientation (Cardin et al., 2007) and frequency (Moore and Wehr, 2013). Similarly, lateral inhibition is mediated by SOM cells in the primary auditory cortex as well (Kato et al., 2017). Based on this, we speculated that auditory gratings would elicit narrowband oscillations. But, contrary to our expectations, auditory gratings induced a broadband response. A recent study has pointed out that the reduced PV^+^ inhibition might enhance broadband high-gamma power due to asynchronous activity (Guyon et al., 2021). So, perhaps auditory gratings activated the cortical neurons but failed to do so synchronously. Differences could also be due to species, as rodent cortex has been shown to generate narrowband gamma (as discussed above). Finally, it could be due to the recording modality. Since the primary auditory cortex is buried in the Heschl’s gyrus, its contribution to the EEG signal may be relatively minor and masked by brain regions on the surface, which may have more complex responses to these auditory gratings.

### Stimulus tuning to gamma and high-gamma responses

We observed a weak gamma tuning in EEG for visual and auditory gratings, though gamma shows a strong tuning in the LFP (Ray and Maunsell, 2010; Jia et al., 2011). MEG studies with visual stimuli have also shown a relatively stronger tuning of gamma oscillations (Adjamian et al., 2004; Koelewijn et al., 2011). Even fMRI studies using auditory gratings demonstrated that voxel activation and transfer functions show tuning towards specific spectro-temporal features of these sounds (Langers et al., 2003; Schönwiesner and Zatorre, 2009). Weaker tuning in EEG could be due to volume conduction effects, which affect other recording modalities (such as MEG) to a lesser degree. We showed strong stimulus tuning in LFP and weak tuning in EEG simultaneously recorded from the same monkeys, which rules out other confounds, such as task and species differences (Murty et al., 2018).

Although we failed to get narrowband gamma using auditory gratings, we found that these stimuli produce very strong and focal broadband high-gamma in EEG. Broadband high-gamma may also prove to be a useful biomarker for diagnosing disorders (Bragin et al., 2010) and a valuable tool for building brain-computer interfaces (Bouchard and Chang, 2014). In addition, differences in the responses between visual and auditory cortices to stimuli that share many similarities may help understand potential differences in their neural circuitry.

## Author Contributions

D.G and S.R. conceived the idea of research and designed the experiments, D.G. collected the data, D.G. and S.R. analyzed the data, and D.G. and S.R. wrote the paper

## Acknowledgements

This work was supported by Wellcome Trust/DBT India Alliance (Senior fellowship IA/S/18/2/504003 to S.R.) and DBT-IISc Partnership Programme. D.G. is thankful for the support from the senior research fellowship awarded by the Council of Scientific and Industrial Research (CSIR).

